# Seasonal gene-expression signatures of delayed fertilization in Fagaceae

**DOI:** 10.1101/2023.03.02.530775

**Authors:** Akiko Satake, Kayoko Ohta, Noriko Takeda-Kamiya, Kiminori Toyooka, Junko Kusumi

**Affiliations:** Department of Biology, Faculty of Science, Kyushu University, Fukuoka 819-0395, Japan; Technology Platform Division, Mass Spectrometry and Microscopy Unit, RIKEN Center for Sustainable Resource Science, Yokohama 230-0045, Japan; Department of Environmental Changes, Faculty of Social and Cultural Studies, Kyushu University, Fukuoka 819-0395, Japan

## Abstract

In the family Fagaceae, fertilization is delayed by several weeks to one year after pollination, leading to one- or two-year fruiting species depending on whether fruiting occurs in the same or the next year after flowering. To investigate physiological responses underlying the regulation of delayed fertilization, we monitored seasonal changes in genome-wide gene expression in tissues including leaves and buds over two years under natural conditions in one- (*Quercus glauca*) and two-year fruiting species (*Lithocarpus edulis*). Genes associated with the responses to cold stress, photosynthesis, and cell proliferation, which are essential for survival and growth, showed highly conserved seasonal expression profiles regardless of species. However, seasonal expression profiles diverged between the one- and two-year fruiting species in genes associated with pollination, an important process contributing to the origin and maintenance of the reproductive barrier between plant species. By comparing seasonal progression of ovule development and gene expression in pistillate flowers, we revealed that ovules started developing after winter in the two-year fruiting species, which could be linked to the activation of genes involved in fertilization and female gametophyte development after winter. These findings suggest that the two-year fruiting species may have evolved a requirement of winter cold to prevent fertilization before winter and facilitate fertilization and embryo development in the following spring when temperature rises. This study offers new possibilities to explore the evolution of reproductive strategies in Fagaceae.

## 1 Introduction

Successful fertilization is the start of a new individual in sexual reproduction. In flowering plants, pollen grains (male gametophytes) arriving on the stigma germinate, and pollen tubes grow to fertilize the egg cell of the female gametophyte generally within 24 to 48 h or even less (Williams, 2008). However, a delay in fertilization more than 4 days was recorded over a century ago (Benson, 1894) and has since been reported in diverse taxa, including Fagales, Brassicales, Laurales and others (Sogo & Tobe, 2006). The family Fagaceae, the most diverse tree family in northern temperate regions, including oaks and beeches, contains an exceptional number of species with delayed fertilization (Satake & Kelly, 2021). A lapse in time between pollination and fertilization spans from several weeks to almost one year. Species in the genus *Fagus* fertilize their ovules five weeks after pollination (Sogo & Tobe, 2006), resulting in fruiting in the same year as flowering (one-year fruiting). In the genus *Lithocarpus*, 92% of 104 species ripen their fruits in the year after pollination (two-year fruiting; Satake & Kelly, 2021). The genus *Quercus* comprises a mixture of one- and two-year fruiting species (Satake & Kelly, 2021).

Satake & Kelly (2021) presented a hypothesis that explains the coevolution of flowering/fruiting phenology and delayed fertilization. They developed a mathematical model that takes into account the impact of winter seasons, which are unfavorable for reproduction, as well as competition for pollinators. By incorporating data on reproductive phenology, they were able to explore how these factors influence the evolution of fertilization timing. When flowering occurs late in the season, particularly during summer or fall, it can be challenging to achieve complete seed maturation before the onset of the cold winter season for Fagaceae species that produce large acorns. The mathematical model has predicted that a strategy of delaying fertilization until after the winter season could evolve as a way to overcome this challenge (Satake and Kelly 2021). The theoretical prediction suggests that the appropriate response to seasonal environmental changes, particularly the response to cold winter, is important for adjusting the fertilization time and aligning it with other phenological traits such as flowering and fruiting time. However, little is known about what physiological responses underlie the regulation of fertilization timing and how they differ between one-year and two-year fruiting species.

The physiological responses to seasonal environmental changes can be studied at the molecular level using the molecular phenology approach (Kudoh, 2016), which monitors the seasonal dynamics of global gene expression profiles in leaves and buds under natural conditions. Recent technological advances in the field of genomics have made it possible to obtain time-course transcriptome data in natural settings in non-model organisms such as Fagaceae (Satake et al. 2022). Molecular phenology has been used to unravel gene expression patterns that govern plant phenology in a wide range of species, including herbs (Aikawa et al. 2010; Nagano et al., 2019, 2012; Richards et al., 2012; Satake et al., 2013), trees in temperate areas (Cronn et al., 2017; Jokipii-Lukkari et al., 2018; Lu, Gordon, Amarasinghe, & Strauss, 2020; Miyazaki et al., 2014; Satake, Kawatsu, Teshima, Kabeya, & Han, 2019), and trees in the tropics (Kobayashi et al., 2013; Yeoh et al., 2017).

To investigate gene-expression signature of delayed fertilization, here we present the comparative molecular phenology over two years in one- and two-year fruiting species of the Fagaceae family. The one-year fruiting species is *Q. glauca* that starts blooming in April and fruits in the autumn in the same year as anthesis (Fig. 1a). The two-year fruiting species, *L. edulis,* begins flowering mainly in June with a minor flowering event in fall (Fig. S1) and fruits in the next year after flowering (Fig. 1b). Two genera, *Quercus* and *Lithocarpus*, diverged during the Paleocene (Manos & Stanford, 2001; Zhou et al., 2022) and diversified in North America and Asia, respectively. We show that performing comparative transcriptomics using these two genera in the same natural habitat is a powerful approach to identify evolutionary conservation and divergence of physiological responses to environmental changes.

**FIGURE 1.**
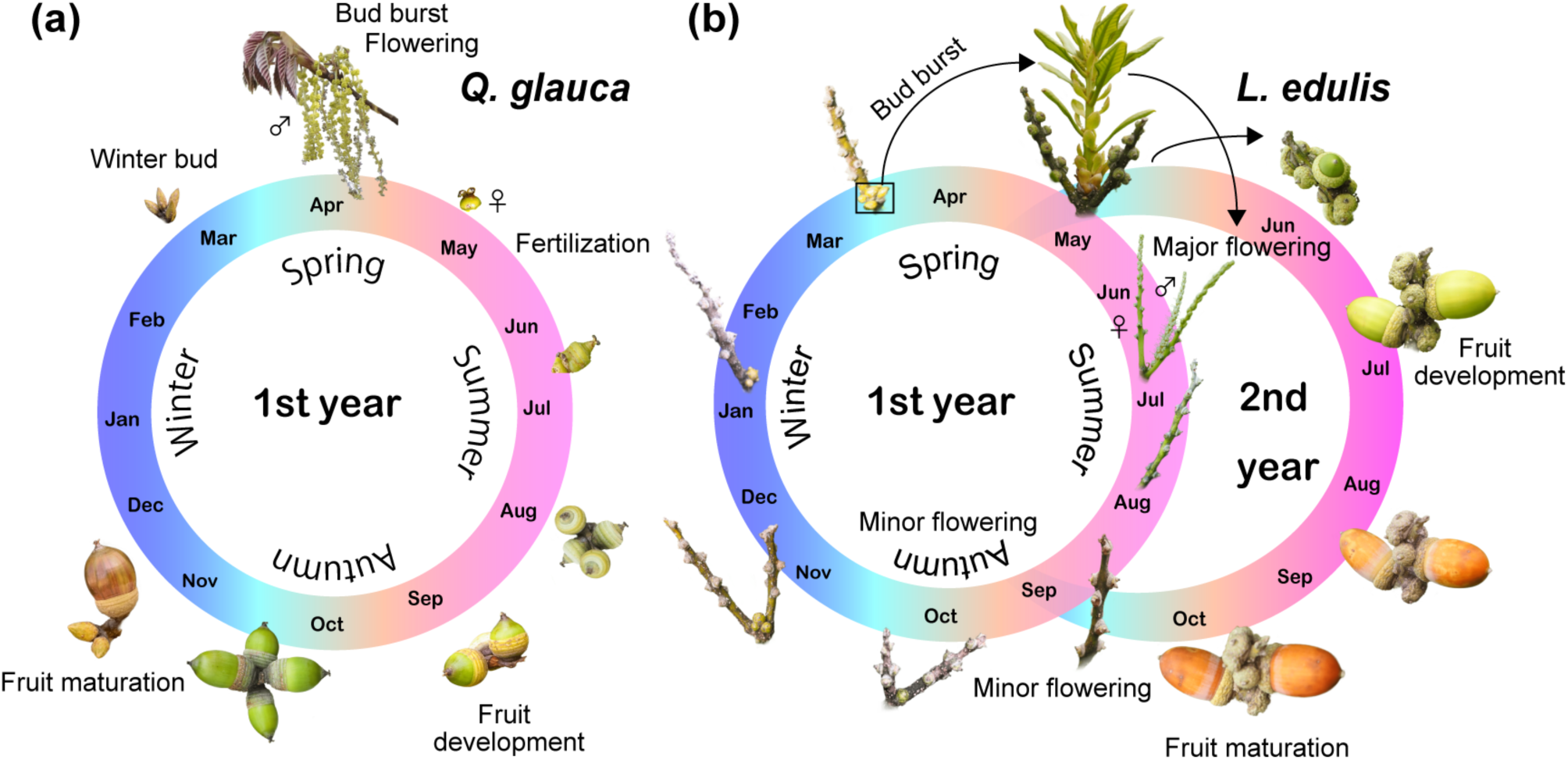
Comparison of flowering and fruiting phenology of *Q. glauca* and *L. edulis*. (a) A one-year fruiting species, *Q. glauca*, starts blooming in April and fruits in the autumn in the same year as anthesis. (b) A two-year fruiting species, *L. edulis*, begins flowering mainly in June with a minor flowering event in fall and fruits in the next year after flowering.

## 2 Materials and Methods

### 2.1 Study site and sampling methods

The study site is in the biodiversity reserve of the Ito campus of Kyushu University (33°35′ 47.5′′ N, 130°12′ 50.0′′ E) situated in Fukuoka, southern Japan. The biodiversity reserve of the Ito campus occupies an area of approximately 37 ha at an elevation from 20 to 57 m a.s.l. The mean annual precipitation and temperature near the site are 1701.0 mm and 16.4 °C, respectively (Japan Meteorological Agency http://www.jma.go.jp/jma/indexe.html for 1991–2020). We studied two evergreen species, *Q. glauca* and *L. edulis*, which are endemic to Asia. The flowers are self- incompatible and wind-pollinated in *Q. glauca,* while they are animal pollinated in *L. edulis*.

We first characterized the seasonal dynamics of global transcriptomics every four weeks using a mixture of leaf and bud tissues over two years because molecular phenology data in floral tissues are available for only several months in one-year fruiting species, which is too short to compare the seasonal progression of gene expression between species. To monitor the molecular phenology, we collected samples from three individuals each for *Q. glauca* (Q1, Q2, and Q3) and *L. edulis* (L1, L2, and L3; Fig. S2). The mean (± s.d.) diameter at breast height (DBH) of *Q. glauca* and *L. edulis* were 17.53 (±1.10) cm and 10.04 (±1.51) cm, respectively. We collected a pair of leaf and apical bud from three current-year shoots per individual every four weeks from May 2017 to February 2019 (sampling dates are provided in Table S1). The pistillate flowers were sampled from three branches per individual from June 2020 to June 2021 in *L. edulis* and from April 2021 to Jun 2021 in *Q. glauca* (Table S1). Samples were taken from the sun-exposed crown (approximately 4 m and 2 m from the ground for *Q. glauca* and *L. edulis*, respectively) using long pruning shears from 11:30 to 12:30 h. For each sample, 0.2–0.4 g of tissue was preserved in a 2 ml microtube containing 1.5 ml of RNA-stabilizing reagent (RNAlater; Ambion, Austin, TX, USA) immediately after harvesting. The samples were transferred to the laboratory within 3 hr after sampling, stored at 4 °C overnight and then stored at −80 °C until RNA extraction. During transport to the laboratory, the samples were kept in a cooler box with ice to maintain a low temperature.

### 2.2 RNA extraction

Total RNA was extracted in accordance with the method described in a previous study (Miyazaki et al., 2014). RNA was extracted independently from leaf and bud samples of each tree and pooled at each time point. Similarly, RNA was extracted independently from each pistillate flower sample. RNA integrity was examined using the Agilent RNA 6000 Nano kit on a 2100 Bioanalyzer (Agilent Technologies), and the RNA yield was determined using a NanoDrop ND-2000 spectrophotometer (Thermo Fisher Scientific). The RNA integrity number (RIN) is listed in Table S2.

### 2.3 Generation of transcriptome next-generation sequencing (NGS) data

We obtained transcriptome data from our samples to design DNA microarray probes. We used eight samples collected monthly from one individual at the study site from May to December 2017 for *Q. glauca* and from June to December 2017 for *L. edulis* (Table S1). Five to six micrograms of total RNA (RNA mixed with equal amounts of RNA extracted from each of the leaves and buds of the same tree) was sent to Macrogen (South Korea), where a cDNA library was prepared with an Illumina TruSeq Sample Prep Kit, and paired-end transcriptome sequencing of each sample was conducted using an Illumina HiSeq2000 or NovaSeq6000 sequencer (Illumina, San Diego, CA, USA). A total of 299 and 313 million 100-bp paired-end reads were obtained for each species. De novo transcriptome assembly was conducted using Trinity version 2.0.6 (Grabherr et al., 2011). Read quality analysis was performed on the raw data using FastQC v0.11.7 (http://bioinformatics.babraham.ac.uk/projects/fastqc/). Quality trimming and adapter clipping were performed using Trimmomatic version 0.38 (Bolger, Lohse, & Usadel, 2014) to trim trailing bases below the average quality 15 and minimum length 36 and clip Illumina adapters. The resulting reads shorter than 50 bp were discarded. De novo transcriptome assembly was conducted using Trinity version 2.0.6 (Grabherr et al., 2011).

### 2.4 Probe design for the DNA microarray

For the custom microarray slides, we used the assembled sequences of the transcripts generated by the NGS described above. We selected the assembled sequences for array design based on two steps. We first extracted transcript sequences that showed high homology against *A. thaliana* (%Identity >= 40%, qcovhsp >= 40%) by BLASTX searches for each species. For each extracted transcript sequence, the top hit *A. thaliana* gene ID was selected. If multiple transcript sequences were annotated for the same *A. thaliana* gene ID, the longest transcript was selected. We obtained 19,290 and 19,426 transcript sequences for *Q. glauca* and *L. edulis*, respectively. In the second step, to select transcript sequences that are conserved across genera in the Fagaceae family, we extracted transcript sequences that were eliminated from the homology selection, but the sequence homology to *F. crenata* transcript sequences used for DNA microarray (Satake et al., 2019) was high (%Identity >= 60%, qcovhsp >= 60%, e-value cut-off: 10^−5^) in BLASTX searches for each species. After the selection in step 2, we obtained additional 3,474 and 4,357 transcript sequences for *Q. glauca* and *L. edulis*, respectively. We pooled these transcript sequences for each species and designed the array using the e-array portal for array design hosted by Agilent (https://earray.chem.agilent.com/earray/) based on 22,764 and 23,783 transcript sequences for *Q. glauca* and *L. edulis*, respectively. Two probes were designed for each transcript sequence. After removing probes with redundant sequence, 42,121 and 42,436 probes were installed in the 8×60K array format.

### 2.5 Microarray analysis

One hundred nanograms of total RNA extracted from the leaves and buds of each sample was amplified, labelled, and hybridized to a 60K Agilent 60-mer oligomicroarray in accordance with the manufacturer’s instructions for each sample for each time point based on the one*-*colour method. The hybridized microarray slides were scanned by an Agilent scanner. The relative hybridization intensities and background hybridization values were calculated using Agilent Feature Extraction Software (9.5.1.1). Among the two probes designed for each transcript sequence, we selected the probe with the largest median. Finally, we obtained time-series data of 15,451 and 15,182 independent probes for *Q. glauca* and *L. edulis*, respectively.

### 2.6 Prediction of orthologous genes

To identify orthologous genes across *Q. glauca* and *L. edulis*, we first used TransDecoder (http://transdecoder.sourceforge.net/) to detect coding regions from the RNA seq assembled contigs. To maximize the sensitivity to capture coding regions with functional significance, we scanned all coding regions detected by TransDecoder for the blastp or pfam searches. We used the protein sequence database of green plants (Viridiplantae) for homology searches with an e-value < 10^−5^. Among the assembled contigs of *Q. glauca* and *L. edulis*, TransDecorder identified 101,371 and 86,128 contigs containing candidate coding regions with homology to known proteins. The longest predicted protein sequences of candidate coding regions were used for subsequent analysis. The construction of groups of orthologous genes (orthogroups, referred to here as gene families including orthologue pairs) was performed for five plant species: *Q. glauca*, *L. edulis*, *Fagus crenata* (75,926 sequences reported in Satake et al. 2019) and *Q. robur* (25,808 sequences from OAK GENOME SEQUENCING http://www.oakgenome.fr), and *Arabidopsis thaliana* (48,359 sequences from TAIR https://www.arabidopsis.org).

The prediction of orthogroups was based on a blastp all-against-all comparison of the protein sequences (e-value < 10^−5^) of these species, followed by clustering with OrthoFinder (Emms & Kelly, 2015, 2019). We obtained 32,149 orthogroups in total and then considered a pair of probes of the two species (*Q. glauca* and *L. edulis*) for which sequences belonged to an identical orthogroup to be orthologous genes. However, certain pairs of probes could not be assigned to be orthologous due to multiple partners within an orthogroup. Additionally, pairs of probes that belonged to an orthogroup lacking the sequence of *A. thaliana* were excluded due to the uncertainty of their function. Finally, we obtained 9,258 pairs of probes predicted to be orthologous genes (Table S3). The GO terms of predicted proteins (orthogroups) were retrieved from annotation data of *A. thaliana*. We removed probes with low signal and weak correlation between individuals using the following three criteria: (1) no signal over all time points, (2) the mean signal value over all time points is lower than 0.05, and (3) the mean of correlation between each pair of individuals is smaller than 0.2. A total of 7,707 pairs among the 9,258 pairs satisfied the criteria. We used time series data of these 7,707 probes for further analyses after normalization to a mean of zero and a standard deviation of one for each species.

### 2.7 Hierarchical clustering

To assess the similarity of the genome-wide transcriptional profiles across orthologous genes, we performed hierarchical clustering using the monthly time series data of 7,707 orthologous genes from March 2017 to February 2019. For each orthologous gene, there were 24 time points, with three individuals each, for *Q. glauca* and *L. edulis*. We calculated the mean expression levels of each orthologous gene across three individuals in each species and subsequently normalized the values by adjusting the mean to zero and the standard deviation to one. We performed hierarchical clustering using the Ward method and the Euclidian distance using the hclust function in R (ver. 3.6.1).

### 2.8 Principal component analysis (PCA)

To assess the seasonal expression dynamics of 7,707 orthologous genes, we performed PCA of the gene expression profiles from all samples using the function prcomp of the stats package in R (ver 3.6.1). To investigate the genes and functions that most contribute to each principal component, we extracted the top 2.5% of genes from the largest positive (*n* = 176) and negative loading values (*n* = 176) for each axis. Then, to test the enrichment of GO terms in each principal component, we performed Fisher’s exact tests (two-sided) using the fisher.test function in R (ver 3.6.1). After the Fisher’s exact tests, we controlled for the false discovery rate using Storey’s *Q*-value method (Storey, 2002) and estimated the *Q*-value of each test using the qvalue package in R (ver 3.6.1). To test the significance of the PCA loadings, we used the bootstrapped eigenvector method (Jackson, 1995; Peres-Neto et al., 2003). Using this method, we confirmed that all the top 2.5% of genes with positive or negative loading values that characterize each axis are considered significant contributors.

### 2.9 Gene ontology (GO) enrichment analysis

To inspect functions of genes for each cluster or those in the top 2.5% of genes that contribute to each of the three principal components, GO enrichment analysis was performed. The 7,707 orthologous genes were selected as a customized reference for the analysis. The list of GO terms for describing the biological process were retrieved from the Database for Annotation, Visualization, and Integrated Discovery (DAVID) (Dennis et al., 2003). Statistical tests for enrichment were performed using Fisher’s exact tests (fisher.test function) in R (ver 3.6.1). We controlled for the false discovery rate using Storey’s *Q*-value method (Storey, 2002) and estimated the *Q*-value of each test using the qvalue package in R (ver 3.6.1). The GO terms with the top-5 lowest *P* value were selected for representation (Supplementary Tables 4 and 5).

### 2.10 Phylogenetic analysis

Based on GO enrichment analyses, we identified three candidate genes, *Secretary* (*SEC*)*5A* (AT1G76850), *SEC15B* (AT4G02350) and *RNA-dependent RNA polymerase 6* (*RDR6*: AT3G49500), that may contribute to the regulation of delayed fertilization. The phylogenetic tree for each of three genes was reconstructed based on the protein sequences of *A. thaliana*, *Oryza sativa*, three *Quercus* species, *L. edulis* and *F. crenata* with *Physcomitrella patens* as an outgroup. Sequences of *SEC5A*, *SEC15B*, and *RDR6* genes for *O. sativa*, *Q. suber,* and *P. patens* were obtained from OrthoDB release 10 (https://www.orthodb.org/) (Kriventseva et al., 2019). Database searches were conducted using annotation keywords or identifiers of the TAIR database. The ortholog sequences of each gene were aligned using muscle implemented in MEGA X (Kumar et al., 2016). Maximum likelihood (ML) trees were constructed with 1,000 replicates for bootstrapping using RAxML v8.2.11 (Stamatakis, 2014) via raxmlGUI 2.0.10 platform (Edler, Klein, Antonelli, & Silvestro, 2021). For each ML estimation, the best substitution model was used, as determined by model testing conducted with ModelTest-NG version 0.1.7 (Darriba et al., 2020) via the raxmlGUI 2.0.10 (Edler et al., 2021). The resulting ML trees were visualized using Figtree version 1.44 (http://tree.bio.ed.ac.uk/software/figtree/).

### 2.11 RT‒qPCR

Pistillate flower samples collected from *L. edulis* from June 2020 to June 2021 were used for RT‒ qPCR analysis. Because pistillate flowers are fertilized in a relatively short period in *Q. glauca*, we used pistillate flower samples from April 2021 to June 2021 for RT‒qPCR analysis in *Q. glauca*. We quantified the expression levels of *SEC5A*, *SEC15B*, and *RDR6* using the expression level of *UBQ10* as a reference and the Bio-Rad CFX connect real-time PCR detection system (CRX96 Touch). cDNA synthesis was carried out using PrimeScript^TM^ RT regent kit with gDNA Eraser (Takara, Japan) from 250 ng total RNA. The first strand cDNAs were diluted to 10 times for subsequent use. Gene specific real-time PCR was performed using 5 ng of cDNA and SsoFast™ EvaGreen® Supermix kit (Bio-Rad, Hercules, CA, USA) according to the manufacturer’s instructions. The PCR condition was as follows: 95 ℃ for 2 min, followed by 40 cycles of 95 ℃ for 1 s, 60 ℃ for 5 s, and the fluorescent signal was measured at the extension step. Melt curve analyses was carried out to validate the specificity of the PCR amplicons. All test cDNAs were run in duplicates for each gene, and the average was used for analysis. The relative expression level of *UBQ10* was normalized to the standard sample by using the comparative threshold cycle (ΔΔCt) method (Pfaffl, 2001). Gene specific primers were designed using Primer3 software (http://primer3.wi.mit.edu/), and confirmed by the observation of a single amplification product of the expected size and sequences. The PCR amplification efficiencies ranged from 102% for to 112% with *R*^2^ > 0.98. At each time point, there are three biological replicates and two technical replicates for each species. We used *UBQ10* as a housekeeping gene because previous studies showed that the expression of *UBQ10* is stable in *F. crenata* (Miyazaki et al., 2014; Miyazaki & Satake, 2017). The mean (± s.d.) Ct values of *QgUBQ10* and *LeUBQ10* in pistillate flowers were 22.0 (± 0.51) and 24.75 (± 0.77), respectively. The primers used for RT‒qPCR are listed in Table S4.

### 2.12 Histological analysis of ovule development

Pistillate flowers in different stages of development were fixed with 4% (w/v) formaldehyde, 2% (v/v) glutaraldehyde, and 0.05 M cacodylate buffer, dehydrated in an ethanol dilution series, and embedded in LR-white resin (Electron Microscopy Science). Sections (1-μm thick) were prepared with an ultramicrotome (Leica Microsystems EM-UC7) using a diamond knife (DIATOME Histo). Each section was stained with 0.05% (w/v) toluidine blue and observed with an upright light microscope with 20x objective lens (Olympus BX53 M and DP26 digital camera).

## 3 Results

### 3.1 Winter and summer transcriptional profiles are distinctly different, but spring and fall are similar

Our global transcriptomic data show clear seasonal dynamics (Fig. 2a). Most orthologues show highly correlated seasonal expression profiles between species (mean of Pearson correlation = 0.39) compared with those calculated from a set of randomly paired genes (mean of Pearson correlation = 0.0096; *P* < 10^-16^; Wilcoxon test; Fig. S3). Hierarchical clustering of the monthly expression profiles revealed clustering of winter months (December, January, February, March) and other seasons regardless of species (Fig. 2a). The transcriptional profiles other than winter were divided into summer months (June, July, August, and September) and spring (April, May, June) or fall (October and November) (Fig. 2a). The expression profiles were similar between spring and fall regardless of the different pheno-phases (bud burst and flowering in spring and fruiting in fall).

**FIGURE 2.**
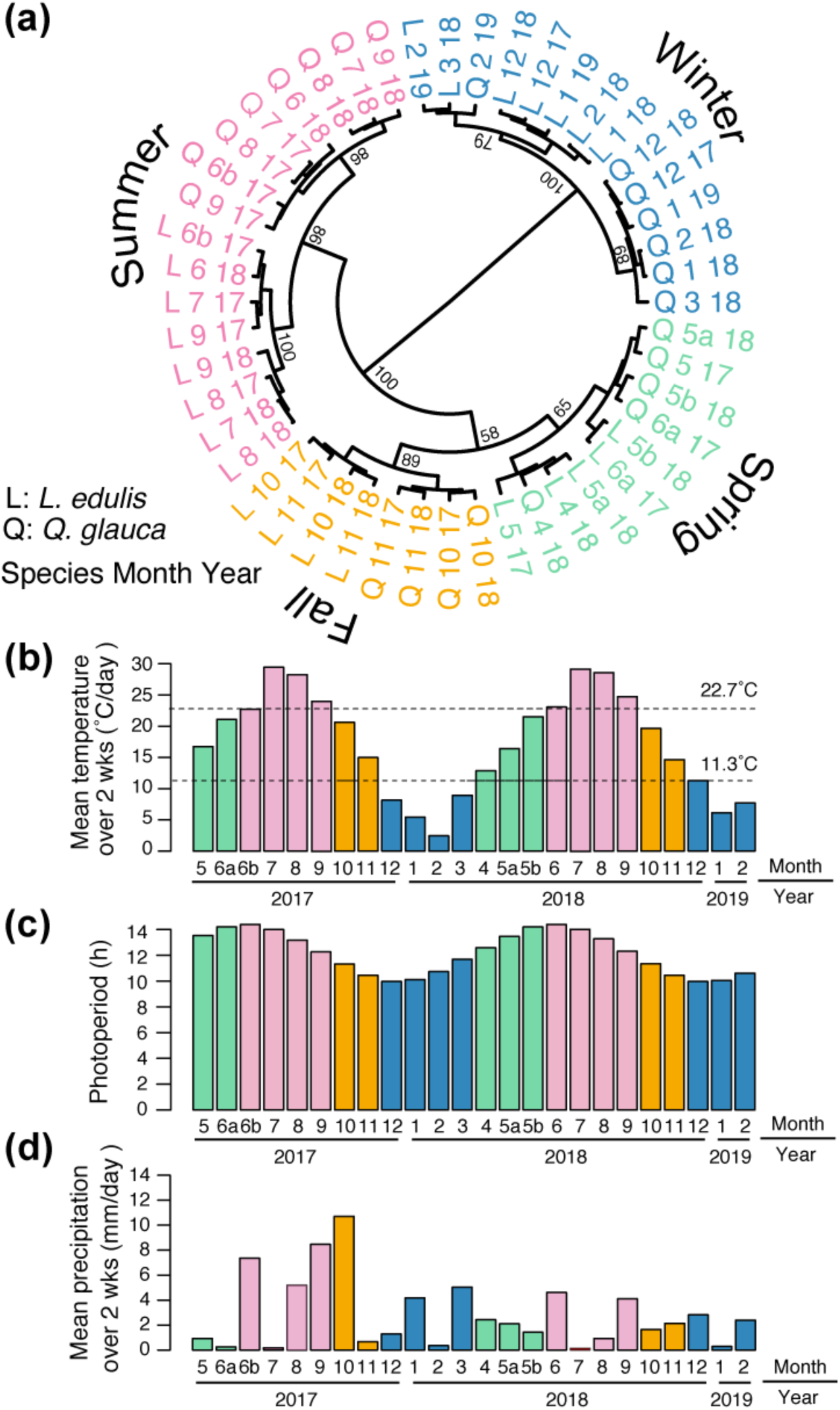
Molecular phenology and seasonal environmental changes. (a) Hierarchical clustering of monthly transcriptome profiles of tissues including leaves and buds in *Q. glauca* (Q) and *L. edulis* (L). The numbers indicate the month and year when each sample was collected. When sampling was performed twice a year, the month is distinguished by a or b (Table S1). The number given to each branch represents the bootstrap confidence level from 1,000 bootstrap samples. (b) The mean temperature over two weeks before the sampling dates. Dashed lines represent 22.7 °C, above which temperature the summer profiles in gene expression emerge, and 11.3 °C, below which temperature the winter profiles in gene expression emerge. (c) The photoperiod on sampling dates. (d) The mean precipitation over two weeks before the sampling dates. The colours in Panels (b), (c), and (d) indicate the gene expression profiles for winter (blue), spring (green), summer (pink), and fall (orange), which are consistent with those in Panel (a).

### 3.2 Genome-wide molecular phenology is associated with seasonal change in temperature, not for photoperiod and rainfall

We found a clear relationship between the transcriptional profiles and seasonal temperature change. A winter transcriptional profile was observed when the mean temperature over the 2 weeks prior to the monitoring date was lower than 11.3 °C (Fig. 2b), and summer transcriptional profiles appeared when the mean temperature exceeded 22.7 °C. Between these temperature thresholds, spring and fall transcriptional profiles were observed (Fig. 2b). The photoperiod and precipitation were not well associated with seasonal transcriptional profiles (Fig. 2c, d). The photoperiod cannot be used to distinguish the differences in transcriptional profiles between spring and summer or those between fall and winter (Fig. 2c). Precipitation fluctuates heavily across months and years, which is different from the seasonal transcriptional profiles (Fig. 2d). These results suggest that temperature is a major driver of seasonal transcriptional dynamics.

Hierarchical clustering of the seasonal expression of individual genes showed that 1,942 genes were highly expressed only in winter (called winter genes) in both *L. edulis* and *Q. glauca* (Cluster 1 in Fig. 3). The biological function of winter genes was enriched by Gene Ontology (GO) terms associated with metabolism (Table S5), implying metabolic changes in response to winter cold. A total of 1,566 genes were highly or moderately expressed in spring and fall (called spring-fall genes) in both species (Cluster 2 in Fig. 3), in which GO terms associated with the cell cycle and cell division were enriched (Table S5), suggesting active cell proliferation in spring or fall seasons. The majority of genes (2,117 genes) were highly expressed in summer (called summer genes) in both species (Cluster 3 in Fig. 3). The top GO term was “Oxidation‒Reduction Process”, suggesting the response to oxidative stress caused by high light and drought in summer (Table S5). The remaining genes, which accounted for 27% of all orthologous genes, showed differential expression between species (Fig. 3). A total of 1,004 genes were expressed in winter in *Q. glauca* but expressed in spring and summer in *L. edulis* (called type 1 differentially expressed genes (DEGs); Cluster 4 in Fig. 3), while 1,078 genes were expressed in winter in *L. edulis* but expressed in spring and summer in *Q. glauca* (called type 2 DEGs; Cluster 5 in Fig. 3). The top five GO terms for type 1 and 2 DEGs included “Oxidation‒Reduction Process” and “Pollination”, respectively (Table S5).

**FIGURE 3.**
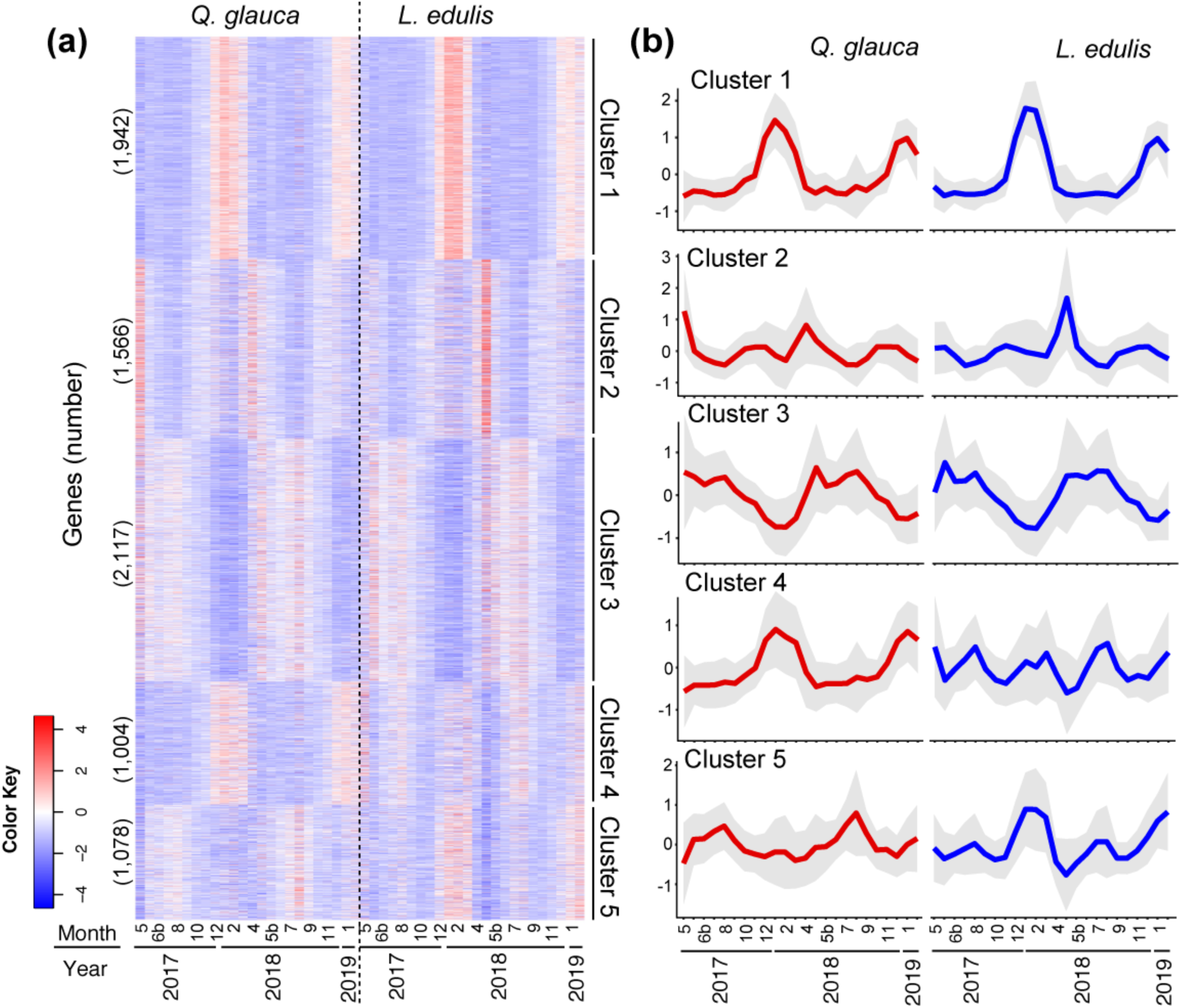
Five representative clusters for seasonal expression profiles of 7,707 orthologues. (a) A hierarchically clustered heatmap of seasonal expression profiles of 7,707 orthologues in tissues including leaves and buds. The numbers in brackets indicate the number of genes included in each cluster. (b) Seasonal expression profiles of genes in each of the five clusters. The average (line) ± s.d. (envelope) is shown (*n* = 1,942, 1,556, 2,117, 1,004, and 1,078 for Cluster 1–5, respectively).

### 3.3 Seasonal gene expression profiles are conserved for the cold stress response, energy acquisition and cell proliferation but diverge for pollination across the two species

To identify and quantify the major axes of the seasonal gene expression profiles, we performed PCA. We found that the first three axes of variation (the principal components; PCs) explain 41.3% of the multidimensional functional space variation (Fig. 4a, b; Fig. S4). The first axis (PC1) accounted for 24.32% of the variation and distinguished gene expression profiles between winter and other seasons (Fig. 4a, c). The second axis (PC2) accounted for 10.58% of the variation and separated gene expression profiles between spring/fall and summer (Fig. 4a, d). Together, the first and second axes represent conserved gene expression dynamics across the two species. In contrast, the third axis (PC3) accounted for 6.45% of the variation and captured the differential gene expression dynamics across the two species (Fig. 4b, e).

**FIGURE 4.**
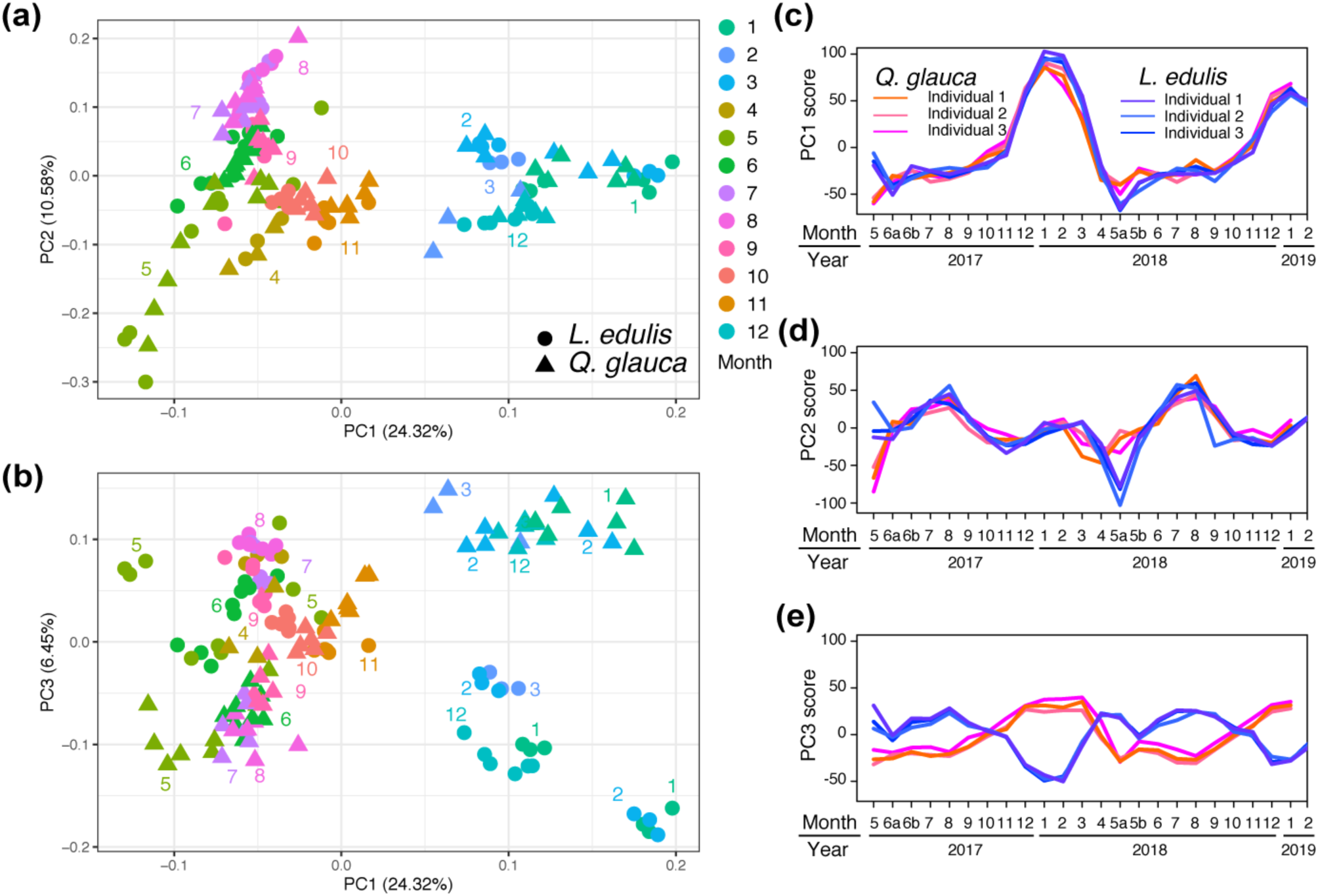
Major axes of multivariate molecular phenology in *Q. glauca* and *L. edulis*. Plot of PC2 versus PC1 (**a**) and PC3 versus PC1 (**b**) resulting from the PCA of the 7,707 orthologues for three individuals per species (triangle: *Q. glauca*, circle: *L. edulis*). The numbers indicate the sampling month. The numbers in brackets represent the explained variance. Plot of the PC1 score (**c**), PC2 score (**d**), and PC3 score (**e**) against the month. Expression profiles were monitored in tissues including leaves and buds.

To identify the key genes responsible for defining each of the three axes, we selected the top 2.5% of genes (*n* = 176 for each axis) with the largest positive and negative loading values. This selection was based on a validation of the significance of each gene’s contribution to each axis. The biological functions of the top 2.5% of genes that characterized the first axis with positive loading were enriched by the GO term “Macromolecule Metabolic Process” (Table S6), which is consistent with the GO term identified in the winter cluster Table S5). Genes associated with cold adaptation (e.g., *Arabidopsis thaliana DEAD-box RNA helicase*: *AtRH7:* AT5G62190) (Huang et al., 2016; Liu et al., 2016), freezing tolerance (e.g., *STARCH EXCESS 1*: *SEX1*: AT1G10760) (Yano et al., 2005; Yu et al., 2001), and proteome and RNA homeostasis (e.g., *Clp protease ATP-binding subunit*: *CLPC1*: AT5G50920) (Nishimura & Van Wijk, 2015; Sjögren, MacDonald, Sutinen, & Clarke, 2004) are included in this GO term (Fig. 5; Table S7), which suggests that the first axis is functionally characterized by the stress response to winter cold. The biological functions of the top 2.5% of genes that characterized the second axis with positive or negative loading were enriched in several GO terms associated with photosynthesis and cell division (Tables S6, S7). The genes associated with photosynthesis were highly expressed in summer (e.g., *STN7*: AT1G68830) (Bellaflore et al., 2005), and those associated with cell division showed an expression peak in spring (e.g., *A. thaliana RADiation54*: *AtRAD54*: AT3G19210 and *METHYLTRANSFERASE1: MET1*: AT5G49160; Fig. 5) (Kankel et al., 2003; Osakabe et al., 2006). This result suggests that the second axis is characterized by energy acquisition and growth. Increased expression of DNA repair and methylation in spring can be the response to DNA replication stress during leaf flushing and growth (Fig. 1). The slight delay in the expression peaks of *MET1* and *AtRAD54* in *L. edulis* (Fig. 5) can be explained by the delayed timing of leaf flushing in *L. edulis*.

**FIGURE 5.**
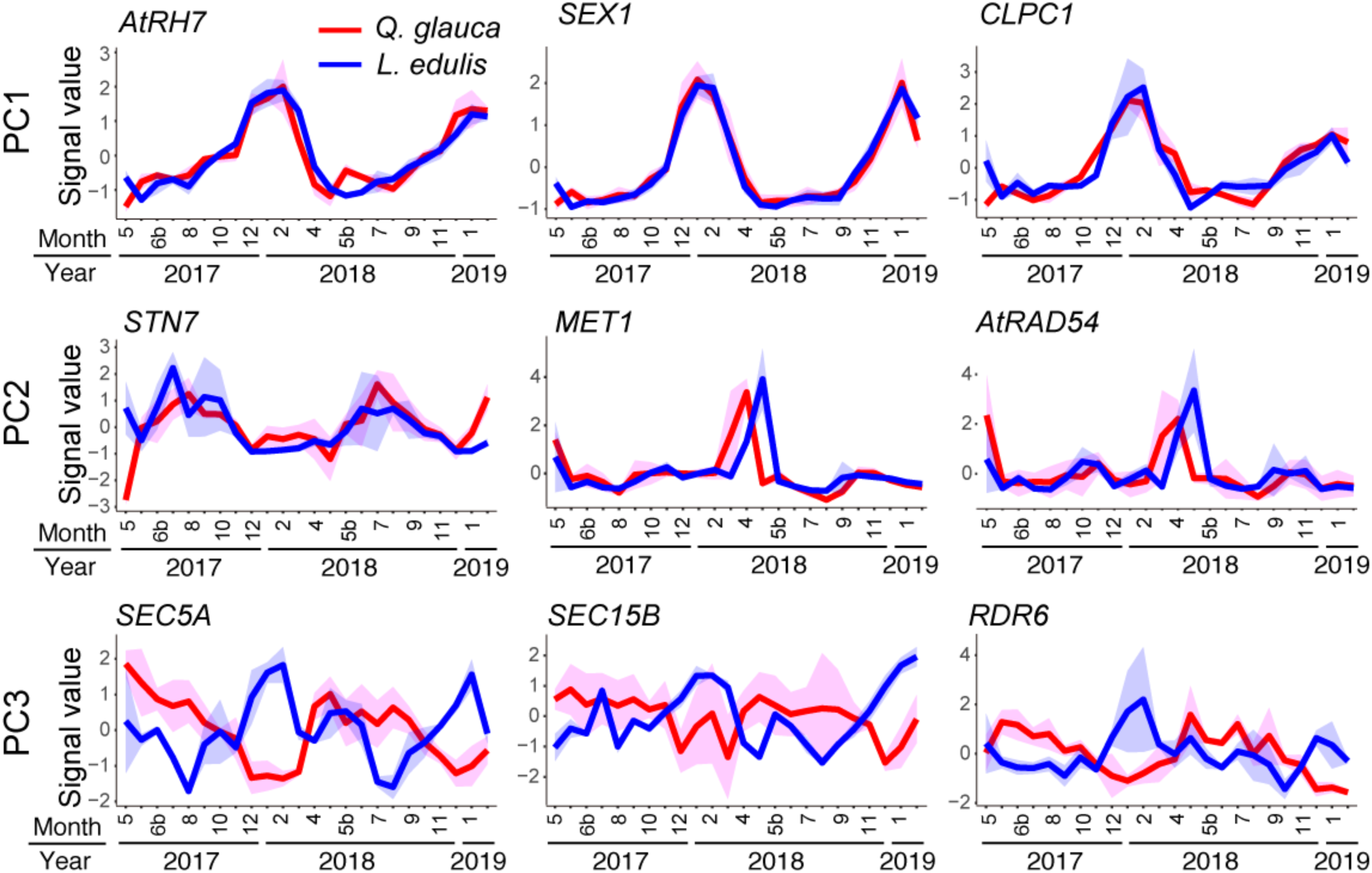
Seasonal expression profiles of genes that characterize each of the three axes. Three genes were selected for each of the three axes as examples: PC1 (*AtRH7, SEX1, and CLPC1*), PC2 (*STN7*, *MET1*, and *AtRAD54*), and PC3 (*SEC5A, SEC15B,* and *RDR6*). The two lines indicate signal values for *Q. glauca* (red) and *L. edulis* (blue). The average (line) ± s.d. (envelope) is shown (*n* = 3). Expression profiles were monitored in tissues including leaves and buds.

The enriched GO terms for the top 2.5% of genes for the third axis with negative loading include “Cell Communications” and “Pollination” (Table S6), consistent with the GO term identified as Type 2 DEGs (Table S5). Because genes included in the GO term “Pollination” could be associated with delayed fertilization, we conducted further analyses on these genes. All eight genes included in the GO term “Pollination” were found to overlap with the genes identified as Type 2 DEGs (Tables S5, S7). Among the eight genes, four genes encode membrane associated proteins, Niemann-Pick type C protein (NPC1-2: AT4G38350), G-type lectin S-receptor-like protein kinase (AT2G19130), and receptor-like protein kinase2 (RPK2: AT3G02130), and signal peptide peptidase (SPP: AT2G03120), involved in sphingolipid trafficking (Feldman, Poirier, & Lange, 2015), anther development (Mizuno et al., 2007), protein secretion (Han, Green, & Schnell, 2009) (Table S7). Purple acid phosphatase 15 (PAP15: AT3G07130) that is an acidic phosphatase with phytase activity (Kuang, Chan, Yeung, & Lim, 2009) was also included. The other three genes that are known to be associated with regulating fertilization on the female side in *A. thaliana.* Among the three genes, two are associated with exocytosis (*SEC5A* and *SEC15B*: Fig. 5). These genes encode subunits of the exocyst, an evolutionarily conserved heterooligomeric protein complex (Heider & Munson, 2012). The exocyst complex tethers and docks vesicles to target membranes (Guo et al., 1999). In *A. thaliana*, through an RNAi approach, these subunits of the exocyst complex were shown to be required for conferring stigma receptivity, suggesting the role of exocytosis as a crucial process during pollen acceptance (Safavian et al., 2015). The last gene is *RDR6* (Fig. 5), which functions in the biogenesis of trans-acting small interfering RNAs (ta-siRNAs) (Peragine, Yoshikawa, Wu, Albrecht, & Poethig, 2004; Yoshikawa, Peragine, Mee, & Poethig, 2005). Loss-of-function mutations in these genes exhibit an increased frequency of abnormal gamete precursors that often give rise to development of more than one female gametophyte in the Arabidopsis ovule (Olmedo-Monfil et al., 2010). Because the association with fertilization has been most thoroughly investigated only in these three genes, we selected *SEC5A*, *SEC15B*, and *RDR6* for further analysis. Because differential seasonal expression profiles of these genes could be associated with the difference in the period of delay from pollination to fertilization, we observed the ovule development by confocal microscopy and compared the ovule development with the expression of *SEC5A*, *SEC15B*, and *RDR6* in pistillate flowers. Before the analysis using pistillate flowers, we confirmed that the three genes of *Q. glauca* and *L. edulis* are found in the clade of corresponding genes of other species in the molecular phylogenetic trees (Figs. S5–S7).

### 3.4 Ovules start developing after winter in two-year fruiting species

The comparative anatomical investigation revealed that *L. edulis* takes 11 months to develop ovules, which is six times longer than *Q. glauca*. In the pistillate flowers of *Q. glauca*, the locules are already visible in May, which is one month after flowering and pollination (Fig. 6a). Six ovule primordia were present, and two dome-shaped ovule primordia arose in each locule (Fig. 6a). These ovule primordia expanded to fill the locules, and one megaspore mother cell was visible in the transverse section in June (Fig. 6a). Because embryo development was observed in July (Fig. S8), fertilization occurred between June and July in *Q. glauca*. In contrast, the pistillate flowers of *L. edulis* were relatively unchanged over eight months after pollination. The three locules were visible, but the ovule primordia were not yet differentiated in the transverse section in November (Fig. 6b). After winter, two dome-shaped ovule primordia arose in each of three locules in March (Fig. 6b). It took an additional two months for the ovules to be well differentiated with megaspore mother cells (Fig. 6b; Fig. S9). In June, one ovule was successfully fertilized, and the zygote enlarged (Fig. 6b). The rest of the ovules showed signs of abortion, as inner integuments and internal structures coagulated to form amorphous, dark-staining structures contained within outer integuments (Fig. 6b).

**FIGURE 6.**
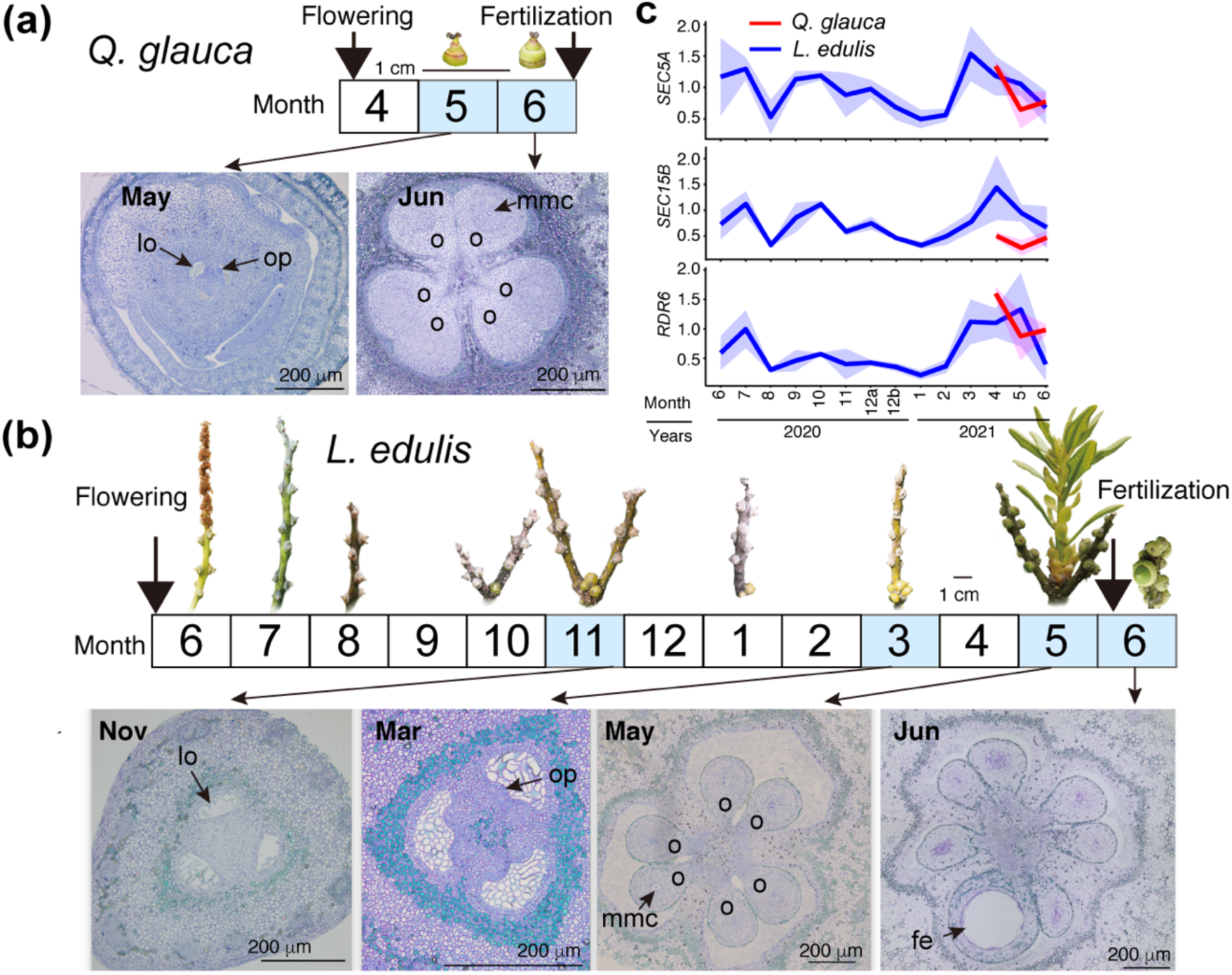
Ovule development and expression dynamics of candidate genes. Transverse sections of ovules and image of pistillate flowers of *Q. glauca* (**a**) and *L. edulis* (**b**) collected on November 11 in 2018 and March 10, May 23, June 6 in 2019. lo: locule, op: ovule primordia, mms: megaspore mother cell, o: ovule, fe: fertilized egg. **c**, Relative expression levels of *SEC5A, SEC15B,* and *RDR6* (average ± s.d. of three replicates) for *Q. glauca* (red) and *L. edulis* (blue) during 2020–2021 against *UBQ10* as a housekeeping gene.

### 3.5 Expression of *SEC5A*, *SEC15B*, and *RDR6* in pistillate flowers increases after winter in two-year fruiting species

We compared the seasonal progression of ovule development with the expression profiles of *SEC5A*, *SEC15b*, and *RDR6* in pistillate flowers quantified by RT‒qPCR. The expression of these genes in *L. edulis* peaked from March to May after winter (Fig. 6c), which is several months later than the peak of leaf and bud tissues (Fig. 5) and coincides with the onset of ovule primordia development (Fig. 6b). The expression of *SEC5A* and *SEC15B* in *L. edulis* showed two other minor peaks in July and October (Fig. 6c), suggesting the intermittent activation of exocytosis. The expression of *SEC5A* and *RDR6* in *Q. glauca* was already high immediately after flowering in April and rapidly decreased in May and June (Fig. 6c). In contrast, the expression level of *SEC15b* was low throughout the census period (Fig. 6c), suggesting that *SEC15b* may be less important for fertilization in *Q. glauca*. Overall, the consistency of expression dynamics and the onset of ovule development suggest that *SEC5A* and *RDR6* are candidates for delayed fertilization in *L. edulis*.

## 4 Discussion

Our results demonstrate that a seasonal gene-expression signature of delayed fertilization is the activation of genes involved in fertilization and ovule development in response to low temperatures. This finding suggests that the two-year fruiting species may have evolved a requirement of winter cold to prevent fertilization before winter and allow for fertilization and embryo development in the following spring when temperature rise.

Our study also revealed evolutionary conserved mechanisms that facilitate appropriate physiological responses to seasonal environmental changes. Specifically, genes associated with the responses to cold stress, photosynthesis, and cell proliferation, which are essential for survival and growth, show highly conserved seasonal expression profiles regardless of species. This indicates that comparative transcriptomics in natural settings is a powerful approach for identifying evolutionary conservation and divergence of physiological responses to environmental changes.

Genes that exhibit divergent expression profiles between one- and two-year fruiting species could potentially be the candidate genes for delayed fertilization. In this study, our focus was on two genes, *SEC5A*, which encodes a subunit of the exocyst complex, and *RDR6*, which is required for posttranscriptional silencing. To validate the requirement of winter cold as an adaptive strategy for adjusting fertilization time, it would be useful to conduct female flower specific transcriptomics to see if their seasonal gene expression profiles are conserved in the two-year fruiting species and identify additional candidate genes of delayed fertilization.

The exocyst complex plays a critical role during pollen‒stigma interactions by mediating the delivery of Golgi-derived secretory vesicles through vesicular trafficking, tethering, and fusion with the plasma membrane for secretion (Cvrčková et al., 2012; Heider & Munson, 2012). The cargo of these secretory vesicles is unknown, but candidate cargo could be plasma membrane aquaporins, which could facilitate water transfer (Safavian & Goring, 2013; Windari et al., 2021) from the stigmatic papilla to the pollen grain for hydration and cell-wall modifying enzymes for stigmatic papillar cell wall loosening and pollen tube penetration (Elleman & Dickinson, 1996; Samuel et al., 2009). In two-year fruiting *Quercus* (*Q. acutissima*, *Q. rubra*, *Q. suber L.*, *Q. velutina*) pollen germinates instantly after pollination to penetrate the stigmatic surface to the stylar transmitting tissue (Cecich, 1997; Deng et al., 2022). Then, the pollen tube is arrested and overwinters at the style-joining site until the formation of the rudimentary ovule and embryo sac maturation in the next spring (Cecich, 1997; Deng et al., 2022). Intermittent and delayed activation of *SEC5A* in the pistillate flowers of *L. edulis* with a sharp expression peak after winter (Fig. 6c) implies female-side regulation for the delivery of secretory vesicles to provide the resources, probably cell-wall modifying enzymes or signalling molecules, necessary for the resumption of pollen tube growth to fertilize the ovule after a prolonged postpollination period. It will be interesting future studies to investigate the pollen tube growth dynamics and exocyst gene expression within the pistil and ovule in various species to elucidate the role of the exocyst complex in male and female gametophyte communication during the long journey from the pollen tube.

Another candidate for delayed fertilization is *RDR6*, which generates ta-siRNAs that essentially silence TEs, which are selfish genetic elements that insert copies of themselves into the genome, during female gametophyte development in Arabidopsis (Olmedo-Monfil et al., 2010). In the pistillate flower of *L. edulis*, *RDR6* was highly expressed after winter, which was synchronized with the timing of female gamete formation and the expression peak of *SEC5A*. These results show that female gamete formation and pollen‒pistil interactions are coordinately regulated in response to seasonal environmental conditions, particularly the winter cold. ta-siRNAs are mobile signal molecules that move from somatic cells into adjacent germline cells (Long et al., 2021; Martínez et al., & Slotkin, 2016; Wu & Zheng, 2019) and even from host plants to fungal pathogens to induce cross-species RNA interference (Cai et al., 2019; Cai et al., 2018). Because exocysts may participate in tethering these extracellular vesicles to the plasma membrane (Saeed et al., 2019), it is tempting to speculate that ta-siRNAs are actively delivered to silence TEs during gametogenesis to protect the genome of the female gamete or enhance male and female gametophyte communication, probably for the selection of compatible pollen tubes in Fagaceae with a high degree of self-incompatibility. To gain a better understanding of how mobile siRNAs function in ovules of Fagaceae and to further illuminate the roles of siRNAs in delayed fertilization, functional analyses and in-situ hybridization experiments would be valuable strategies.

The long delay in fertilization has long been a mystery in biology. This study offers new possibilities to explore the evolution of this unique reproductive strategies.

## Acknowledgements

The authors would like to thank Y. Sawasaki and M. Seki for their supports in sample collection, M. Seki and E. Sasaki for their technical supports with the statistical analysis, and K. Miwa for her valuable comments on our initial draft. We also thank three reviews for their valuable suggestions to improve our manuscript.

## Data Accessibility and Benefit-Sharing

The sequence data and DNA microarray data that support the findings of this study are available from the NCBI BioProject (accession PRJNA872835 for *Q. glauca* and PRJNA872836 for *L. edulis*), the NCBI Shotgun Assembly Sequence Database (TSA) (accession GKBD00000000 for *Q. glauca* and GKBC00000000 for *L. edulis*), and the NCBI GEO database (accession GSE211382, GSE211384, and GSE211385). Benefits from this research accrue from the sharing of our data and results on public databases as described above.

## Author contributions

A.S. conceived of and designed the analysis; K.O. and A.S. collected samples; and K.O. performed the molecular experiments. K.J. and A.S. analysed the data; N.T. and K.T. performed anatomical observation of ovules. A.S. wrote the paper with input from all of the authors. This study was funded by JSPS KAKENHI (JP17H01449, JP21H04781) to A.S.

